# Evaluation of pre-analytical factors affecting plasma DNA analysis

**DOI:** 10.1101/126839

**Authors:** Havell Markus, Tania Contente-Cuomo, Winnie S. Liang, Mitesh J. Borad, Shivan Sivakumar, Simon Gollins, Nhan L. Tran, Harshil D. Dhruv, Michael E. Berens, Alan Bryce, Aleksandar Sekulic, Antoni Ribas, Jeffrey M. Trent, Patricia M. LoRusso, Muhammed Murtaza

## Abstract

Pre-analytical factors can significantly affect circulating cell-free DNA (cfDNA) analysis. However, there are few robust methods to rapidly assess sample quality and the impact of pre-analytical processing. To address this gap and to evaluate effects of DNA extraction methods and blood collection tubes on cfDNA yield and fragment size, we developed a multiplexed droplet digital PCR (ddPCR) assay with 5 short and 4 long amplicons targeting single copy genomic loci (mean amplicon size: 71 bp and 471 bp respectively). Using this assay, we compared performance of 7 cfDNA extraction kits and found cfDNA yield and fragment size varies significantly between them. We also compared 3 blood collection protocols used to collect plasma samples from 23 healthy volunteers (EDTA tubes processed within 1 hour and Cell-free DNA BCT tubes at ambient temperature processed within 24 hours and 72 hours of collection). To assess whether cell-stabilizing preservative in BCT tubes introduced noise in cfDNA, we performed digital targeted sequencing. We found no significant differences in cfDNA yield, fragment size and background sequencing noise between these protocols. In 219 clinical samples tested for quality using the ddPCR assay, cfDNA fragment size was significantly shorter in plasma samples immediately processed for ctDNA analysis compared to archived samples, suggesting background DNA contributed by lysed peripheral blood cells. In summary, we describe a multiplexed ddPCR approach that enables cfDNA quality assessment and could inform the design of future circulating tumor DNA studies.

**Gene names:** None

## Introduction

Circulating cell-free DNA (cfDNA) analysis has found several diagnostic applications in prenatal, transplant and cancer medicine^1-4^. In patients with cancer, cfDNA analysis relies on detection and quantification of somatic alterations to assess tumor-specific cfDNA. Total and tumor-specific cfDNA levels in plasma vary considerably across patients, cancer types and disease stages as well as during longitudinal follow-up of each patient^5,6^. Several recent reports have described sensitive molecular methods for analysis of tumor-specific cfDNA levels^7-9^. However, our understanding of how preanalytical factors affect performance and results of downstream molecular assays is limited^10^.

cfDNA fragments in plasma have a modal fragment size of 160-180 bp, corresponding to DNA protected in mono-nucleosomes^11^. One challenge when analyzing plasma DNA is the variable contribution of high molecular weight (HMW) DNA resulting from lysis of peripheral blood cells during blood processing^12-15^. HMW DNA is not intended to be part of the molecular readout during cfDNA analysis but it can affect PCR and sequencing results. High fractions of HMW DNA in plasma can complicate PCR and tagmentation-based sequencing because these methods will incorporate intact DNA in a sample, potentially biasing the data towards wild-type alleles and increasing false negative results. In contrast, ligation-based library preparation from cfDNA does not require any additional DNA fragmentation and therefore excludes intact DNA. However, if the contribution of intact DNA is not taken into account during sample quantification, library preparation can vary in performance. Ideally, rapid processing of blood samples as soon as possible after venipuncture can overcome these issues but real-time processing of samples is challenging in clinical environments. With growing interest in cfDNA-based diagnostics, several solutions have emerged to streamline pre-analytical processing. These include special blood collection tubes that contain cell-stabilizing preservative to minimize lysis of peripheral blood cells for up to several days after venipuncture. In addition, cfDNA-focused extraction kits have been introduced that claim preferential extraction of fragmented cfDNA over HMW DNA from the same sample.

There is scarcity of robust methods that allow quality assessment of low input cfDNA samples and a comparison between pre-analytical solutions. Here, we present a multiplexed digital PCR approach that can reliably assess cfDNA quantity and contribution of HMW DNA. We use this assay and next-generation sequencing to compare cfDNA extraction kits and blood collection tubes with current gold standards and to perform quality assessment of plasma samples across multiple clinical cohorts.

## Results

### Droplet Digital PCR to assess cfDNA concentration and fragment size

To enable reliable assessment of amplifiable DNA concentration and fragment size using minimal quantities of cfDNA, we designed a multiplexed ddPCR assay targeting 9 single copy genomic loci^16^. We included 5 short PCR amplicons with mean product size of 71 bp (range 67-75 bp) and corresponding probes labeled with FAM as well as 4 long PCR amplicons with mean product size of 471 bp (range 439-522 bp) and corresponding probes labeled with TET (Figure 1 and Supplemental Table 2). We expected two populations of droplets on ddPCR with distinct fluorescence, each representing the sum of products from the two amplicon sets. Assuming no copy number changes, the number of FAM-positive droplets represents 5 times the number of haploid genome equivalents (GEs) with fragments long enough to be amplified using short amplicons. Similarly, the number of TET-positive droplets represents 4 times the GEs with fragment sizes amplifiable using long amplicons. We calculated low molecular weight cfDNA concentration as the difference between short and long GEs.

**Figure 1:**
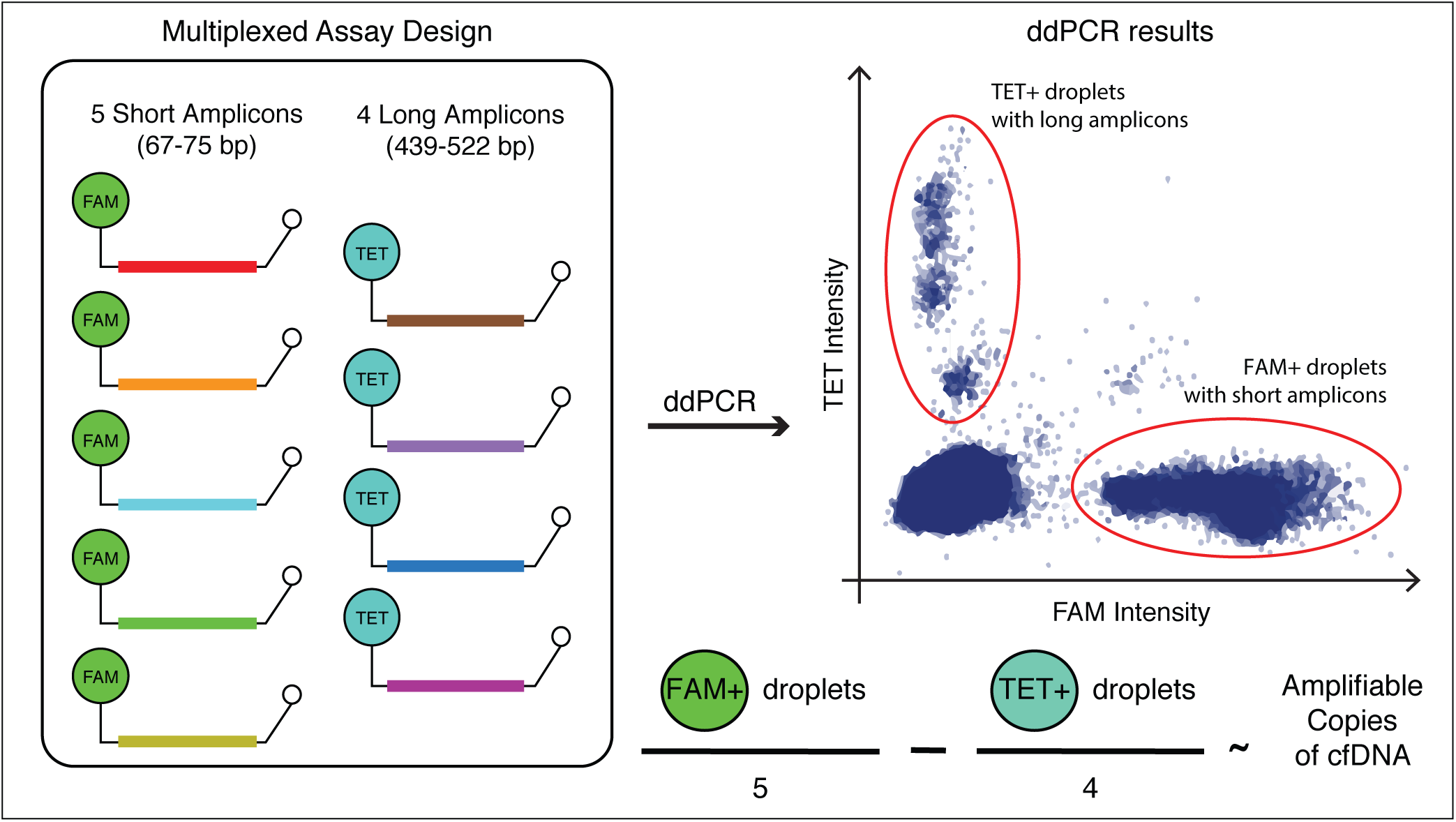
Schematic representation of a multiplexed droplet digital PCR approach for measuring cfDNA quantity and evaluating fragment size.

To evaluate the performance of this approach, we analyzed known quantities of control DNA with predetermined fragment sizes and compared our results with expected PCR performance given amplicon and fragment size. Observed ddPCR results agreed with expected results for 150, 300, 500 and 1,000 bp fragments remarkably. For example, using a 71 bp amplicon and input of 150 bp DNA fragments, we expect to recover ∼62% and we observed 75% recovery using short amplicons in our assay (Figure 2A). To assess whether ddPCR quantification of input DNA can help overcome variability in sequencing results, we measured the concentration of LMW DNA in 5 control plasma samples to prepare and sequence 12 exome libraries made from 1, 2 and 5 ng LMW DNA. Besides input quantity, all other library preparation conditions including number of PCR cycles were similar between these replicates. Library size and diversity was estimated from 0.83-1.66 million read pairs per sample. As expected, library diversity correlated strongly with input DNA quantities (Pearson r=0.938, p=2.48 × 10^−7^; Figure 2B).

**Figure 2:**
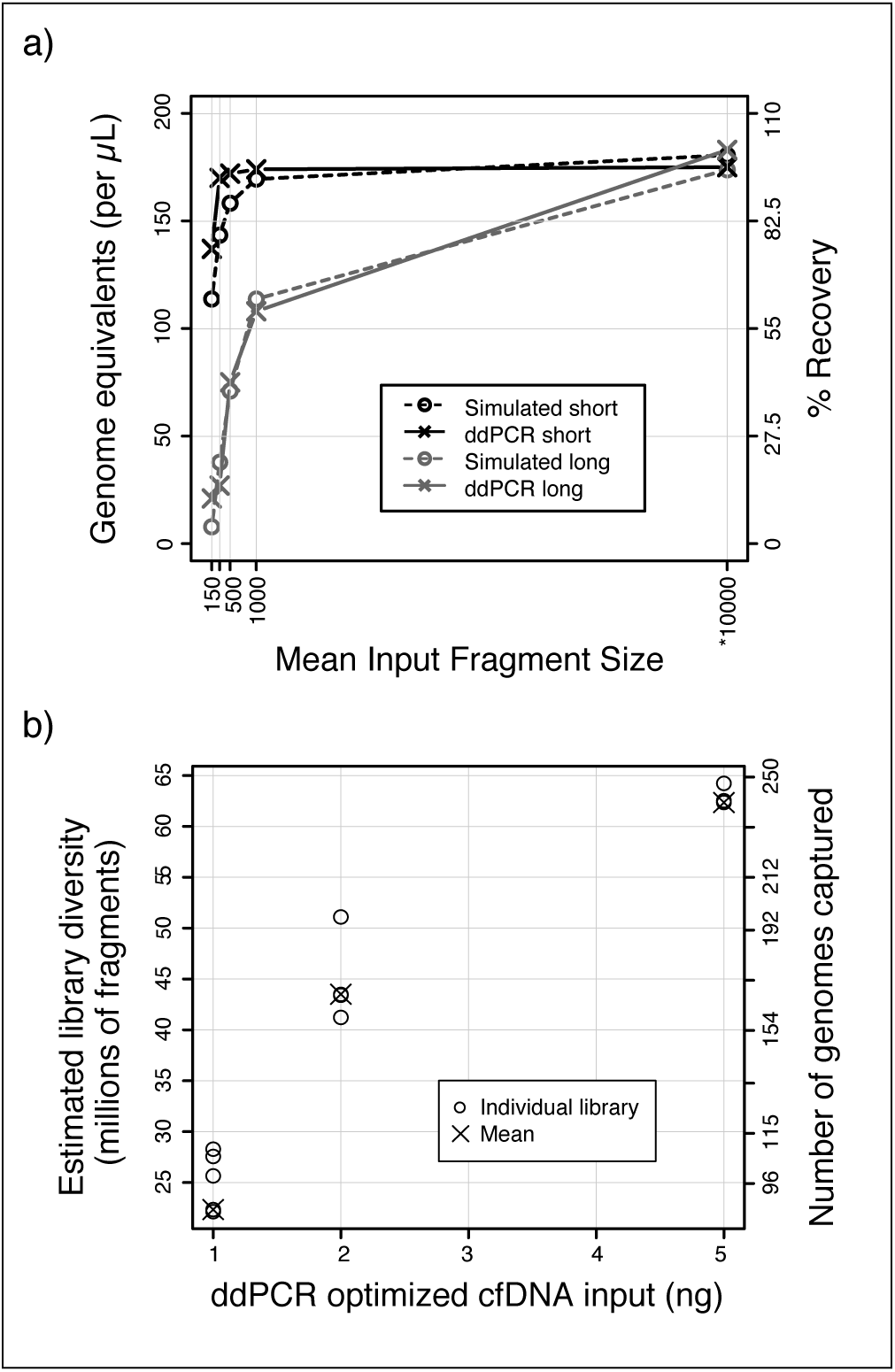
Evaluation of ddPCR assay performance. **a)** Comparison of ddPCR measurements with expected results when using control DNA of known fragment size as template. Solid lines with crosses show measured ddPCR results. Dotted lines with circles show simulated data. *Simulated data was generated assuming 10,000 bp fragments but ddPCR measurement was performed on intact genomic DNA (without sonication). **b)** Evaluation of sequencing library diversity obtained with cfDNA input amounts measured using the ddPCR assay. On the second y-axis (right), library diversity has been converted into number of genome equivalents, assuming 65 Mb exome captured region and 180 bp fragment size. 5, 4, and 3 individual libraries were prepared for 1, 2, and 5 ng cfDNA input respectively.

### Comparison of cfDNA extraction kits

To compare performance of cfDNA extraction kits and minimize the contribution of biological variation, we obtained pooled control plasma sample from a commercial vendor. We compared 7 different extraction kits marketed for cfDNA extraction including 3 spin column-based methods and 4 magnetic beads-based methods (labeled A-G, see Supplemental Table 1). For each kit, we performed 10 replicates of DNA extraction from 1 mL plasma and quantified cfDNA yield and fragment size using ddPCR. We found wide variability in yield and fragment size across these kits (ANOVA p=5.01 × 10^−11^ and p=1.16 × 10^−11^ respectively, Figure 3A and 3B). Highest median yield of LMW cfDNA was obtained using Kit A that uses spin columns for cfDNA extraction (1,936 GEs/mL of plasma, n=10). Median LMW fraction for Kit A was 89%. The median yield of LMW DNA using Kit B was not significantly different than Kit A but the results were more variable (1,760 LMW copies/mL plasma, t-test p=0.427). Amongst methods based on magnetic beads, Kit E showed the highest yield of LMW DNA (median 1,515 LMW copies/mL, n=10) and a LMW fraction comparable to Kit A (median 90%). In comparison to Kit A, the yield was significantly lower (t-test p=9.46 × 10^−5^).

**Figure 3:**
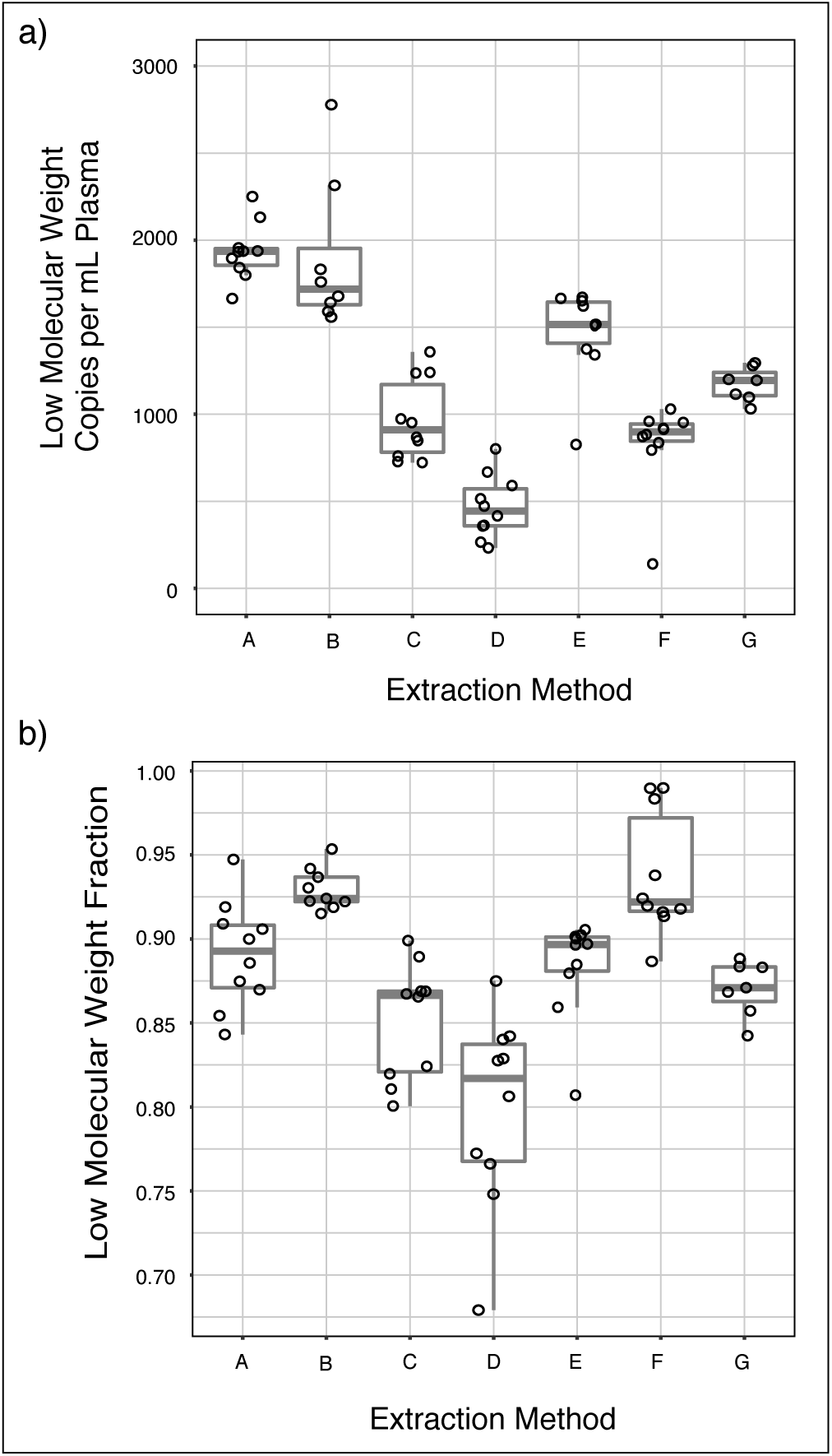
Evaluation of cfDNA extraction methods using ddPCR. Kits labeled A-C use spin columns and kits labeled D-G use magnetic beads for cfDNA isolation. For each extraction kit, n=10 individual extractions were performed successfully, starting with 1 mL plasma volume except for kit B (n=9) and kit G (n=7). Aliquots from a single pool of 250 mL commercially obtained control plasma sample were used for these experiments. **a)** Comparison of cfDNA yield (in LMW copies / mL of plasma) across 7 extraction kits. An outlier at 5,306 LMW copies / mL for Kit B is not shown to enhance scaling. **b)** Comparison of LMW fraction across 7 extraction kits.

### Comparison of blood collection protocols

To investigate how cfDNA yield and LMW fractions were affected by blood collection tubes and protocols, we collected 3 blood samples each from 23 healthy volunteers (12 males and 11 females): one EDTA tube processed within 1 hour and two Cell-free DNA BCT tubes stored at ambient temperature for 24 hours and 72 hours prior to processing. We processed all samples identically and extracted cfDNA using QIAamp Circulating Nucleic Acid kit (QIAGEN). We measured cfDNA yield and fragment size using ddPCR. Mean LMW GEs/mL plasma for EDTA, BCT 24hr, and BCT 72hr were 1,925, 1,592, and 1,516 respectively with no significant difference between collection protocols (ANOVA p=0.440, Figure 4A). LMW fractions for EDTA, BCT 24hr, and BCT 72hr were 87%, 88%, and 90% respectively with no significant difference (ANOVA p=0.074, Figure 4B).

**Figure 4:**
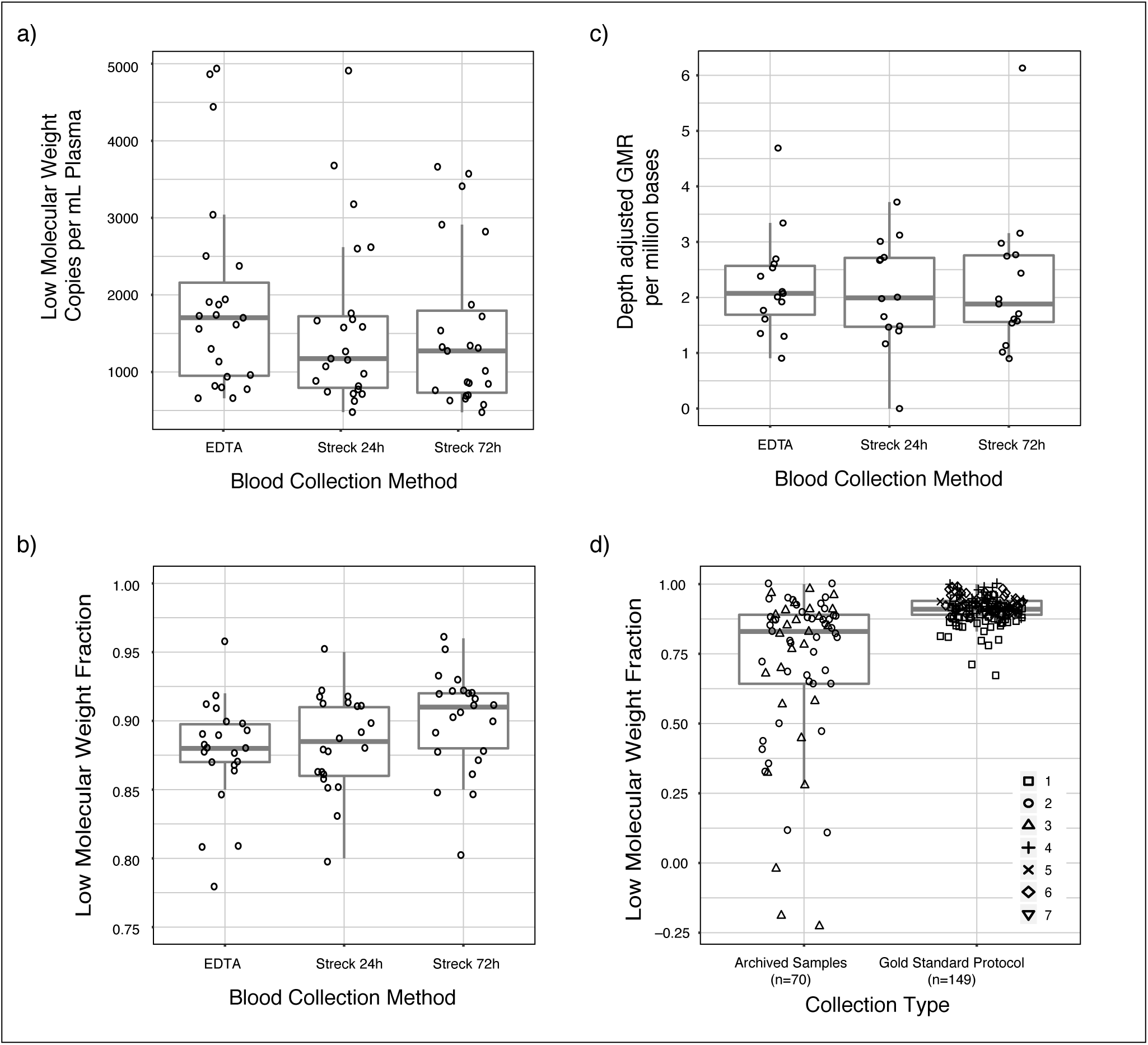
Evaluation of blood collection protocols for cfDNA analysis using ddPCR and digital targeted sequencing. **ab)** Comparison of cfDNA yield (in LMW copies / mL of plasma) and LMW fraction across 3 extraction methods. Outliers at 0.67 LMW fraction for EDTA and at 0.71 LMW fraction for Streck 24 hr are not shown in **b** to enhance scaling. Paired samples from healthy volunteers were collected for this analysis. Comparison between EDTA, Streck 24 hr samples and Streck 72 hr samples showed no significant difference for yield or LMW fraction (ANOVA p-values >0.05). **c)** Comparison of depth-adjusted GMR observed across collection protocols. Pairwise comparisons between EDTA and Streck 24 hr samples (n=13 pairs) and between EDTA and Streck 72 hr samples (n=13) showed no significant difference between collection protocols (paired t-test p>0.05). **d)** Comparison of LMW fraction between archived retrospective samples and samples processed using gold standard protocols for ctDNA analysis, showing significant difference between the two groups (t-test p=2.302 × 10^−7^). Symbols indicate cohort numbers with details in Supplemental Table 3.

To assess whether cell-stabilizing preservative would affect background noise in cfDNA sequencing, we prepared targeted sequencing libraries using a molecular tagging approach that can help distinguish non-reference alleles arising in the original template molecules from errors introduced during PCR amplification. We sequenced 44 samples including all 3 pairs from 12 individuals and 8 additional samples. We generated an average of 8.16 million read pairs per sample and achieved mean unique coverage in the target region of 33.1 read families (with an average 4.26 members/read family). To measure background noise, we calculated global nucleotide mismatch rate (GMR) as the sum of the number of read families with non-reference alleles divided by the sum of unique coverage across all targeted loci (additional details in Methods). GMR was 37.8 ± 12.2, 30.8 ± 9.11 and 30.8 ± 7.77 for EDTA, BCT 24 hr and BCT 72 hr samples respectively (mean ± standard deviation). GMR between paired samples from the same individuals was highly correlated across tube types (Pearson r=0.652, 0.647 and 0.758, p=0.022, 0.017 and 0.003 for the three pairwise comparisons, Supplemental Figure 1). This suggests that biological noise (in vivo) was a much larger contributor to measured GMR instead of noise introduced during sample processing or analysis. Since depth of coverage would determine our ability to measure low-abundance noise, we adjusted GMR by unique depth of coverage and found no significant difference between EDTA and BCT 24 hr as well as EDTA and BCT 72 hr samples (paired t-test p=0.998, n=13 pairs and p=0.451, n=13 pairs respectively, Figure 4C).

### Evaluation of cfDNA fragment size in clinical samples

To evaluate the extent to which background DNA from peripheral cell lysis affects cfDNA analysis in archived retrospective samples, we evaluated a cohort of 219 clinical samples collected across 7 cohorts and multiple collection sites (Supplemental Table 3). Using the ddPCR assay, we found that LMW fractions were significantly lower in archived samples (cohorts 1 and 2, n=70 samples, median LMW fraction: 82.9%) compared to plasma samples collected prospectively for cfDNA analysis and immediately processed using current gold standard protocols (cohorts 3-7, n=149 samples, median LMW fraction: 91.4%, t-test p=2.302 × 10^−7^, Figure 4D).

## Discussion

Variation in pre-analytical processing of plasma samples can affect cfDNA analysis results^12^. Circulating cell-free DNA is predominantly fragmented and delays between venipuncture and isolation of plasma can lead to higher background of intact DNA contributed by lysis of peripheral blood cells^13^. This affects PCR and tagmentation-based sequencing methods by diluting the measured mutation fractions and it affects ligation-based sequencing methods by lowering effective template available for analysis. We have developed a multiplexed droplet digital PCR approach to reliably assess amplifiable quantities of cfDNA and estimate the contribution of high molecular weight background DNA in a single step. As an upfront sample quality assessment assay, this approach can help optimize input quantities to achieve reproducible performance in sequencing experiments. The assay design targets 9 different regions in the genome, lowering minimum input DNA quantity needed for quality assessment. qPCR or digital PCR assays that target 1-2 loci can be biased by locus-specific amplification bias or by somatic copy number changes in plasma samples from advanced cancer patients^17^. Instead, we rely on average readouts across multiple targets to achieve high precision from limited quantities of cfDNA and to accommodate the wide range of total cfDNA concentrations found in patients with cancer (particularly when implemented in ddPCR with millions of partitions).

Several new solutions to streamline cfDNA extraction have been introduced recently including magnetic-bead based approaches. However, most published reports do not include a comparison of these with current gold standards^17-20^. To provide an update and compare cfDNA extraction performance using a robust approach, we evaluated seven commercially available kits marketed for cfDNA extraction and found that QIAamp Circulating Nucleic Acid kit provided the highest cfDNA yield and low molecular weight fractions. Newer methods that use magnetic beads for DNA extraction may be more automatable and in our results, the MagMAX Cell-free DNA Isolation kit provided the highest yield and low molecular weight fractions amongst such workflows.

To overcome the need for rapid processing of plasma samples, special blood collection tubes are available that include a preservative to prevent lysis of peripheral blood cells for up to several days at room temperature^21^. Current applications for cfDNA analysis in prenatal diagnostics rely on relative representation of genomic regions but in patients with cancer, there is need for detection of mutations at very low fractional abundance and preservative induced noise could lead to false positives. In paired samples from healthy volunteers, we found no evidence that cfDNA yield or fragment size was significantly different between EDTA tubes processed within 1 hour of collection and BCT tubes stored at ambient temperature and processed 72 hours after collection. These results are in agreement with published observations^22-25^. In addition, we also extensively explored whether preservative in BCT tubes induced any pre-analytical noise using digital targeted sequencing. To ensure that we could measure pre-analytical noise separately from any errors introduced during library preparation, we used a molecularly tagged sequencing strategy. We found no evidence that global mutation rate (adjusted for depth of sequencing) is any different in blood stored in BCT tubes at ambient temperature for up to 72 hours after collection as compared to samples collected in EDTA tubes and processed within 1 hour. Interestingly, we observed that GMR was highly correlated between independent replicates from the same individual across blood processing protocols, even after thorough filtering for common and private germline SNPs. This suggests that once sequencing errors are filtered out, biological noise introduced in vivo is the predominant contributor to GMR instead of pre-analytical errors introduced in vitro. Such biological noise could represent non-specific somatic mutations (potentially originating in circulating blood cells) or extracellular damage to cfDNA during circulation. Although we cannot distinguish between these mechanisms based on our results, this observation warrants further study particularly as the field investigates circulating tumor DNA analysis for early detection of cancer.

There are multiple limitations of this study that must be acknowledged. Library preparation kits differ in their efficiency and performance and can be affected by multiple factors such as adapter composition, input DNA concentration and ligation time. While the absolute measurements of library yield and diversity may not be valid for other library preparation methods, accurate quantification and size assessment of input cfDNA will still be relevant. Similarly, several factors can affect DNA extraction yield including input plasma volumes and elution volumes. When comparing cfDNA extraction methods, we minimized biological background by testing 70 replicates of 1 mL each from a single pooled plasma sample. Extraction performance of these kits may differ from our results if a different volume of plasma is used. For comparison of blood collection and processing protocols, we performed paired analysis of samples and found no appreciable differences in background noise between EDTA and BCT tubes.

Our effective sample size for the sequencing-based comparison is limited to 13 pairs and that could limit power to detect subtle differences in pre-analytical error rates but there are no indications of even a trend towards a difference in depth-adjusted GMR between collection protocols. In addition, these comparisons were performed on samples obtained from healthy volunteers and we cannot directly comment on whether tumor-specific mutant DNA will behave similarly.

In summary, we have developed a droplet digital PCR based approach to assess quantity and quality of plasma DNA samples to improve performance of downstream sequencing assays. We presented comparison of several methods for cfDNA extraction and blood collection for cfDNA and their potential effects of downstream analysis. We now routinely use this assay for quality assessment of all plasma samples processed and analyzed for ctDNA studies in our lab. We find significant differences in quality between archived samples and prospectively collected plasma samples, processed rapidly for ctDNA analysis, highlighting the importance and need for appropriate preanalytical processing of samples. Since archived samples from clinical trials are often accompanied by long-term follow-up and clinical annotation, their use can be critical to evaluate clinical utility of ctDNA analysis in cancer patients. The ddPCR approach we describe here will be useful to assess quality of archived samples prior to analysis. Our findings can benefit the design of future studies and clinical trials by helping minimize the contribution of pre-analytical variability in circulating tumor DNA analysis.

## Methods

### Droplet Digital PCR to assess quantity and fragment size

We designed a multiplexed ddPCR assay targeting 9 single copy genomic loci^16^. We included 5 short PCR amplicons with mean product size of 71 bp (range 67-75 bp) and corresponding probes labeled with FAM as well as 4 long PCR amplicons with mean product size of 471 bp (range 439-522 bp) and corresponding probes labeled with TET (Figure 1 and Supplemental Table 2). We used PrimerQuest (Integrated DNA Technologies) to design these assays, evaluated them to avoid known polymorphic loci and used in silico PCR to confirm each amplicon yielded a single product and no cross products when used in multiplex.

We prepared digital PCR reactions at 50 μL volume using 25 μL of 2X KAPA PROBE FAST Master Mix (Kapa Biosystems), 2 μL of 5 mM dNTP Mix (Kapa), 2 μL of 25x Droplet Stabilizer (RainDance Technologies), 9 μL of 100 μM primer mix (pooled equimolarly), 1.68 μL of 20 μM each probe (IDT), 0.25 μL molecular biology grade water and 2-10 μL of input DNA. We generated droplets using RainDrop Digital PCR Source Instrument (RainDance), performed thermocycling using DNA Engine Tetrad 2 (Bio-Rad Laboratories) with the following parameters: 1 cycle of 3 min at 95 °C, 50 cycles of 15 sec at 95 °C and 1 min at 60 °C with a 0.5 °C/sec ramp from 95 °C to 60 °C, 1 cycle of 98 °C for 10 min and hold at 4 °C forever. We measured droplet fluorescence using RainDrop Digital PCR Sense Instrument (RainDance) and analyzed results using accompanying software.

Identification of positive droplets requires setting fluorescence thresholds (gates) for each ddPCR assay (Figure 1). We compared results across intact genomic DNA and no template controls to identify thresholds for this assay. We calculated fractional loss of volume (dead volume) as the difference between measured number of intact droplets of expected size (5 pL) and number of expected droplets (10 million droplets for 50 μL reactions). Any reactions with >50% loss of volume were excluded from further analysis.

### Evaluation of assay performance

For any given PCR reaction, yield is affected by amplicon size and template fragment size. To evaluate whether our ddPCR assay performed as expected, we analyzed known quantities of sheared DNA fragments with predetermined size and compared the results with expected theoretical yields. To generate experimental data, we analyzed 4 aliquots of 0.6 ng/μL human genomic DNA (Sigma Aldrich), sheared by sonication to achieve fragment sizes of 150 bp, 300 bp, 500 bp and 1,000 bp. To estimate theoretical yield, we performed simulations of DNA fragmentation as described previously^26^. Assuming a DNA molecule spanning 2,500 bp on either side of the average amplicon size for each set (5,071 bp for short amplicons and 5,471 bp for long amplicons), we generated break points by sampling from an exponential distribution with a rate representing the corresponding fragment size (150, 300, 500 or 1,000 bp). We determined that a molecule will be “missed” by an amplicon if a DNA break fell within the amplicon region (bound by 5’ ends of the two primers). We sampled 50,000 molecules to determine overall frequency of missed molecules for each combination of amplicon size and fragment size.

### Comparison of cfDNA extraction kits

We obtained 250 mL of a control pooled plasma sample collected with K2 EDTA additive from BioreclamationIVT. Upon receipt, the sample was thawed and stored in 1 mL aliquots at -80 °C until further analysis. We compared 7 different extraction kits marketed for cfDNA extraction including 3 spin column-based methods and 4 magnetic beads-based methods (labeled A-G, see Supplemental Table 1). Manufacturer recommended protocols for plasma DNA extraction were followed. Prior to each extraction, plasma aliquots were centrifuged at 14,000 *g* for 10 minutes to remove any cellular debris.

### Comparison of blood collection protocols

To compare cfDNA yield, fragment size and sequencing noise across blood collection protocols, we collected plasma samples from healthy volunteers. Under an approved protocol (Western IRB protocol number 20142638), we collected 8-10 mL blood in K2 EDTA BD Vacutainer tubes and two Cell-Free DNA BCT (Streck) from each volunteer^21^. EDTA tube was processed within 1 hour of collection. cfDNA BCT were stored at ambient temperature and processed 24 and 72 hours after collection. Samples were centrifuged at 820 *g* for 10 min at room temperature. 1 mL aliquots of plasma were further centrifuged at 16,000 *g* for 10 mins to pellet any remaining cellular debris. The supernatant was stored at -80 °C until DNA extraction. DNA was extracted using the QIAamp Circulating Nucleic Acid kit (QIAGEN). cfDNA yield and fragment size were evaluated using ddPCR. Baseline sequencing noise was evaluated using digital sequencing, targeting a panel of recurrent cancer genes.

### Clinical samples

Plasma samples included in comparison of immediately processed and archived samples were collected with informed consent under multiple independent clinical protocols, all approved by the relevant Institutional Review Boards and approved under Western IRB protocol number 20142638. Samples from 7 independent clinical cohorts were included in this analysis, including patients diagnosed with melanoma, cholangiocarcinoma and rectal cancer as well as healthy volunteers described above. Sample processing conditions and number of samples from each cohort are described in Supplemental Table 3. 7 samples, all from the same ddPCR chip, were excluded from this analysis due to >50% loss of volume during ddPCR. Median dead volume across remaining samples was 10.4% (n=219 samples).

### Sequencing library preparation

For assessment of sequencing performance guided by ddPCR results, we quantified cfDNA from control plasma samples (commercially obtained) using ddPCR and prepared whole genome sequencing libraries using ThruPLEX DNA-Seq (Rubicon Genomics), as per manufacturer’s instructions. We compared performance across 1, 2 and 5 ng amounts of input DNA and concentrated samples to achieve 10 μL starting volume if needed (using vacuum concentration). We assigned sample specific barcodes to each library, quantified them using qPCR and pooled them at equimolar concentrations for target enrichment. We performed exome enrichment using NimbleGen SeqCap EZ Human Exome v3 kit (Roche), as per manufacturer’s instructions except the use of xGen Universal Blocking Oligos – TS HT-i5 and TS HT-i7 (IDT). We quantified all enriched exome libraries using qPCR and pooled at equimolar concentrations for sequencing. We performed sequencing on MiSeq (Illumina) to generate 75 bp paired-end reads and a 6 bp barcode read.

For comparison of background noise between blood collection tubes, we prepared whole genome sequencing libraries from 1 ng cfDNA from volunteer plasma samples using ThruPLEX Tag-seq (Rubicon). This kit introduces a 6 bp random molecular tag on both sides of DNA fragments, making it possible to distinguish any non-reference alleles (true mutations or pre-analytical noise) in the original template molecule from PCR noise introduced during library preparation. We quantified the libraries and pooled them for target capture using xGen^®^ Pan-Cancer Panel (IDT), following manufacturer’s instructions.

### Sequencing data analysis

We demultiplexed sequencing data based on sample-specific barcodes and converted to fastq files using Picard tools v2.2.1, allowing 1 bp mismatch and requiring minimum base quality of 30. We aligned sequencing reads to the human genome hg19 using bwa mem v0.6.2^27^. We sorted and indexed bam files using samtools v1.3.1^28^ and calculated library diversity using Picard tools.

For data comparing background noise across blood collection tubes, we added a barcode (BC) tag comprised of the two paired 6 bp unique molecular identifiers (UMIs) to each aligned read. We parsed these BAM files using a custom R script to identify UMI families and assign read groups as follows: For each set of reads that started and ended at the same genomic positions, we sorted UMI pairs by the number of exact duplicates observed. We used the most frequent UMI pair in this set to seed the first read family. For the remaining reads, if either of the 2 UMIs in a pair were within a Hamming distance of 1 from the corresponding seed UMI, we assigned them to the same read family. For any UMIs not assigned in the first cycle, we repeated this process starting with the next most frequent UMI pair until all reads had been assigned read families. We grouped reads from the same family by adding RG tags to the BAM files. To facilitate computational processing, we assumed fragment sizes in our libraries would not exceed 1,000 bp.

Once all reads were assigned RG tags, we generated pileups per read-family using samtools mpileup. Each read family with ≥ 3 members was included in further analysis. We required ≥ 80% of reads supporting a non-reference allele to call a variant. All variants overlapping with dbSNP v147 were removed. To remove any private SNPs, we filtered out positions covered by < 10 read families or total non-reference allele fraction ≥ 20%. We calculated global nucleotide mismatch rate (GMR) as the sum of the number of read families with non-reference alleles divided by the sum of unique coverage across all targeted loci. To account for variability in depth of coverage between samples, we adjusted GMR for average unique coverage in each sample.

## Acknowledgments

We gratefully acknowledge support for this study through a Bisgrove Scholars Award from Science Foundation Arizona. We also acknowledge support from The V Foundation, Stand Up To Cancer, Ben and Catherine Ivy Foundation and Desert Mountain CARE. We would like to thank Stephanie Althoff and Stephanie Buchholtz at TGen, volunteers and patients who participated in this study.

## Author Contributions

MM and TC designed the study. TC, HM and WSL performed the experiments. MJB, SS, SG, NLT, HDD, MEB, AB, AS, AR, JMT and PML designed and performed clinical studies and performed clinical interpretation. HM and MM performed data analysis and wrote the manuscript. All authors reviewed the manuscript.

## References

1 Bianchi, D. W. et al. DNA sequencing versus standard prenatal aneuploidy screening. The New England journal of medicine 370, 799–808, doi: 10.1056/NEJMoa1311037 (2014).

2 De Vlaminck, I. et al. Circulating cell-free DNA enables noninvasive diagnosis of heart transplant rejection. Science translational medicine 6, 241ra277, doi: 10.1126/scitranslmed.3007803 (2014).

3 Haber, D. A. & Velculescu, V. E. Blood-based analyses of cancer: circulating tumor cells and circulating tumor DNA. Cancer Discov 4, 650–661, doi:10.1158/2159-8290.CD-13-1014 (2014).

4 Lo, Y. M. & Chiu, R. W. Genomic analysis of fetal nucleic acids in maternal blood. Annual review of genomics and human genetics 13, 285–306, doi:10.1146/annurev-genom-090711-163806 (2012).

5 Bettegowda, C. et al. Detection of circulating tumor DNA in early- and late-stage human malignancies. Science translational medicine 6, 224ra224, doi: 10.1126/scitranslmed.3007094 (2014).

6 Dawson, S. J. et al. Analysis of circulating tumor DNA to monitor metastatic breast cancer. The New England journal of medicine 368, 1199–1209, doi: 10.1056/NEJMoa1213261 (2013).

7 Forshew, T. et al. Noninvasive identification and monitoring of cancer mutations by targeted deep sequencing of plasma DNA. Science translational medicine 4, 136ra168, doi:10.1126/scitranslmed.3003726 (2012).

8 Newman, A. M. et al. Integrated digital error suppression for improved detection of circulating tumor DNA. Nat Biotechnol 34, 547–555, doi:10.1038/nbt.3520 (2016).

9 Wan, J. C. et al. Liquid biopsies come of age: towards implementation of circulating tumour DNA. Nat Rev Cancer 17, 223–238, doi:10.1038/nrc.2017.7 (2017).

10 Wong, D. et al. Optimizing blood collection, transport and storage conditions for cell free DNA increases access to prenatal testing. Clinical biochemistry 46, 1099–1104, doi: 10.1016/j.clinbiochem.2013.04.023 (2013).

11 Jiang, P. et al. Lengthening and shortening of plasma DNA in hepatocellular carcinoma patients. Proc Natl Acad Sci U S A 112, E1317–1325, doi: 10.1073/pnas.1500076112 (2015).

12 Sherwood, J. L. et al. Optimised Pre-Analytical Methods Improve KRAS Mutation Detection in Circulating Tumour DNA (ctDNA) from Patients with Non-Small Cell Lung Cancer (NSCLC). PLoS One 11, e0150197, doi:10.1371/journal.pone.0150197 (2016).

13 Chiu, R. W. et al. Effects of blood-processing protocols on fetal and total DNA quantification in maternal plasma. Clinical chemistry 47, 1607–1613 (2001).

14 Jung, M., Klotzek, S., Lewandowski, M., Fleischhacker, M. & Jung, K. Changes in concentration of DNA in serum and plasma during storage of blood samples. Clinical chemistry 49, 1028–1029 (2003).

15 Chan, K. C., Yeung, S. W., Lui, W. B., Rainer, T. H. & Lo, Y. M. Effects of preanalytical factors on the molecular size of cell-free DNA in blood. Clinical chemistry 51, 781–784, doi:10.1373/clinchem.2004.046219 (2005).

16 Eisenberg, E. & Levanon, E. Y. Human housekeeping genes, revisited. Trends Genet 29, 569–574, doi:10.1016/j.tig.2013.05.010 (2013).

17 Devonshire, A. S. et al. Towards standardisation of cell-free DNA measurement in plasma: controls for extraction efficiency, fragment size bias and quantification. Anal Bioanal Chem 406, 6499–6512, doi:10.1007/s00216-014-7835-3 (2014).

18 Mauger, F., Dulary, C., Daviaud, C., Deleuze, J. F. & Tost, J. Comprehensive evaluation of methods to isolate, quantify, and characterize circulating cell-free DNA from small volumes of plasma. Anal Bioanal Chem 407, 6873–6878, doi: 10.1007/s00216-015-8846-4 (2015).

19 Repiska, G., Sedlackova, T., Szemes, T., Celec, P. & Minarik, G. Selection of the optimal manual method of cell free fetal DNA isolation from maternal plasma. Clin Chem Lab Med 51, 1185–1189, doi: 10.1515/cclm-2012-0418 (2013).

20 Fleischhacker, M. et al. Methods for isolation of cell-free plasma DNA strongly affect DNA yield. Clin Chim Acta 412, 2085–2088, doi:10.1016/j.cca.2011.07.011 (2011).

21 Fernando, M. R. et al. A new methodology to preserve the original proportion and integrity of cell-free fetal DNA in maternal plasma during sample processing and storage. Prenat Diagn 30, 418–424, doi:10.1002/pd.2484 (2010).

22 Kang, Q. et al. Comparative analysis of circulating tumor DNA stability In K3EDTA, Streck, and CellSave blood collection tubes. Clinical biochemistry 49, 1354–1360, doi:10.1016/j.clinbiochem.2016.03.012 (2016).

23 Toro, P. V. et al. Comparison of cell stabilizing blood collection tubes for circulating plasma tumor DNA. Clinical biochemistry 48, 993–998, doi: 10.1016/j.clinbiochem.2015.07.097 (2015).

24 Parpart-Li, S. et al. The effect of preservative and temperature on the analysis of circulating tumor DNA. Clin Cancer Res, doi:10.1158/1078-0432.CCR-16-1691 (2016).

25 El Messaoudi, S., Rolet, F., Mouliere, F. & Thierry, A. R. Circulating cell free DNA: Preanalytical considerations. Clin Chim Acta 424, 222–230, doi:10.1016/j.cca.2013.05.022 (2013).

26 Xie, J., Crooke, P. S., McKinney, B. A., Soltman, J. & Brandt, S. J. A computational model of quantitative chromatin immunoprecipitation (ChIP) analysis. Cancer Inform 6, 138–146 (2008).

27 Li, H. Aligning sequence reads, clone sequences and assembly contigs with BWA-MEM. arXiv preprint arXiv:1303.3997 (2013).

28 Li, H. et al. The Sequence Alignment/Map format and SAMtools. Bioinformatics 25, 2078–2079, doi:10.1093/bioinformatics/btp352 (2009).

